# Charged nanobubbles in culture media differentially affect viability of human iPSC-derived neurons

**DOI:** 10.1101/2025.08.31.673401

**Authors:** Yifan Liu, Takeshi Ohdaira, Emi Kitakata, Michael A. Silverman, Jhunam Sidhu, Jun Okubo, Yoshihisa Harada, Kumiko Hayashi

## Abstract

Nanobubbles (NBs) are gas-filled spherical structures less than 1 µm in diameter, characterized by high surface charge, long-term stability, and the ability to generate bactericidal hydroxyl radicals upon collapse. Although these unique properties enable their extensive applications, including wastewater treatment, their cell biological applications at neutral pH remain limited by difficulties in maintaining stability and surface charge. Here, we succeeded in generating both positively and negatively charged NBs, achieving high zeta potentials in human iPSC-derived neural progenitor cell (NPC) and neuron culture media (pH 7.4). Similar to their bactericidal effects, NBs induced cell death in iPSC-derived cells, with positively charged NBs showing stronger cytotoxicity than negatively charged ones. This may be because the cell membrane carries a negative charge, making positively charged NBs more likely to approach and interact with the membrane, or because positively charged nanobubbles exhibit stronger radical generation. In future, we aim to elucidate the molecular biological mechanisms underlying NB-induced cell death and to contribute to the potential application of nanobubbles in regenerative medicine. This study will serve as an initial step toward that goal.

## Introduction

Nanobubbles (NBs) are gas-filled, spherical bubbles with diameters of less than 1 µm, defined as ultrafine bubbles under the international standard ISO/TC281 on fine bubble technology^1^. NBs are generated from microbubbles via a hydrodynamic activation method^2^. The concentrated and persistent presence of electrolytic ions in the aqueous solution surrounding a NB forms an inorganic shell that stabilizes the NB, a process known as the salting-out phenomenon^2^. Similar to colloidal particles suspended in water, which are known to be charged, NBs also exhibit surface charges. Although the mechanism by which NBs acquire surface charges remains unclear, it is likely that the clustered structure of water molecules at the gas–liquid interface plays a key role. Even when the concentration of NBs is high, they do not readily coalesce due to electrostatic repulsion between them, which contributes to their stability.^3–5^

NBs typically have a high specific surface area due to their small size and low buoyancy, which causes them to exhibit Brownian motion. Thermodynamic and kinetic analyses both suggest that NBs cannot remain stable for long periods.^6^ In theory, NBs should dissolve rapidly because of their extremely high Laplace pressure, which is generated by surface tension at the gas–liquid interface and increases as bubble size decreases. However, in experiments, microscopy studies have shown that NBs can remain stable for extended periods; for example, total internal reflection fluorescence microscopy (TIRFM) has been used to detect NBs,^7^ while interference-enhanced reflection microscopy (IERM) has been applied to visualize NB surfaces.^8^ A possible reason is that the surface charge of NBs can reduce the required gas concentration inside a bubble to balance the Laplace pressure.

NBs can generate large amounts of hydroxyl radicals upon collapse.^9^ Hydroxyl radicals generated by the bubble collapse exhibit bactericidal activity^1,10,11^. These unique properties are entirely different from those of larger bubbles (microbubbles), and enable their extensive applications,^12^ including wastewater treatment,^1^ medicine, plant cultivation,^13^ and biomedical applications.^14^ In this study, we aim to investigate whether charged NBs can stably persist in cell culture media, and to examine their effects on human induced pluripotent stem cell (iPSC)-derived neural progenitor cells (NPCs) and neurons. Until now, it has been difficult to generate NBs that remain stable for extended periods in neutral media (pH 7.4). Cell culture media are complex liquids containing a variety of low- and high-molecular-weight compounds, including amino acids, and proteins. Although previous studies have generated NBs in Dulbecco’s Modified Eagle Medium (DMEM) for fibroblast cell culture,^14^ they typically disappear within a few days, making it difficult to apply them to cell cultures that require long-term maintenance for differentiation, such as iPSC-derived neurons.

We successfully generated NBs with an extremely high zeta potential (several tens of mV) and incorporated them into cell culture media for human iPSC-derived neural progenitor cells (NPCs) and neurons. Here, the type of gas used for the NBs was air, and NBs were generated directly in the cell culture media, rather than being produced in a separate solution and subsequently mixed into it. In addition to achieving high zeta potentials, we succeeded in producing positively charged NBs, despite the longstanding challenge of generating them in a stable form. While previous studies have reported that NBs typically exhibit a negative charge in media at pH 7.4,^3–5^ we found that positively charged NBs could remain stable in such environments for over a month. To explore these effects, we compared iPSC-derived cell cultures exposed to either positively or negatively charged NBs. Similar to their bactericidal effects attributed to hydroxyl radicals, in the present study, we observed that NBs induced cell death in iPSC-derived cells. By measuring the number of live cells by using fluorescence micrographs, we were able to quantitatively assess differences in the charged properties of NBs. Furthermore, in terms of software, we developed software utilizing an overlapping ROI (region of interest) scanning method to enable accurate cell counting. Our approach enables such quantitative comparisons, allowing for systematic investigation of effects of NBs. To our knowledge, this is the first report demonstrating the long-stable generation of charged NBs in cell culture media for iPSC-derived cells, enabling future studies on applications of regenerative medicine.

## Results

### Physicochemical properties of charged NBs containing cell culture media

The air NB-containing cell culture medium for human iPSC-derived NPCs was prepared as described in the Methods section. NBs were not added to the cell culture medium afterward; instead, we generated NBs directly in the cell culture medium using an activation plate (Methods). Here, the pH of the cell culture medium was measured to be approximately 7.5 (Fig. 1a). Under such near-neutral conditions, only the generation of negatively charged NBs has been previously reported.^3–5^ However, we successfully generated positively charged NBs in the cell culture medium at approximately pH 7.5. As for physicochemical properties other than pH, although the exact composition of the STEMCELL Technologies media is proprietary, typical reference values for commonly used cell culture media (e.g., DMEM) are as follows: ionic strength 150–170 mM; dissolved oxygen concentration under atmospheric conditions (21% O₂) of 180 µM O₂ (0.18 mM); and osmolality of 280–320 mOsm/kg.

**Fig. 1.**
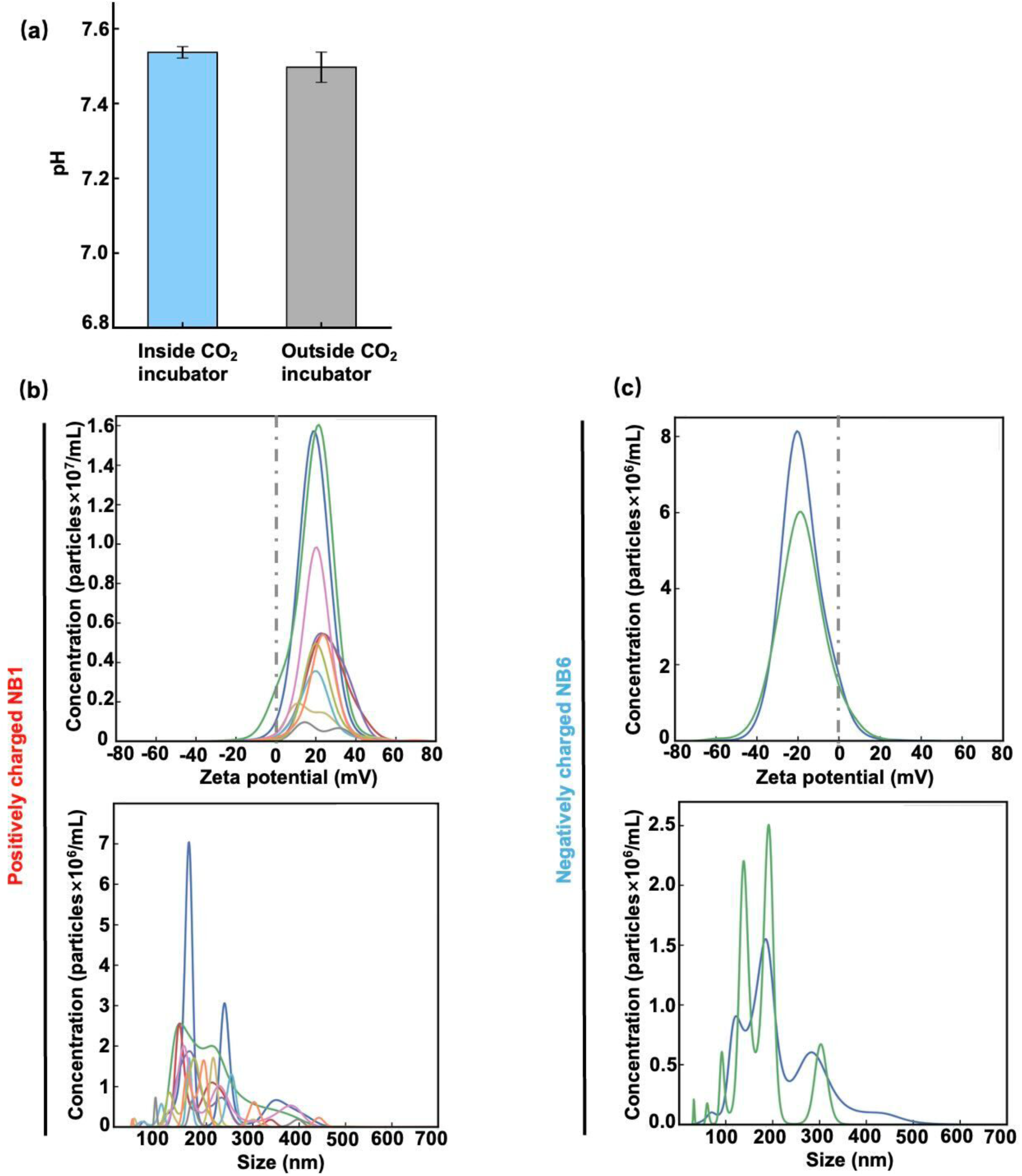
Physical properties of nanobubbles (NBs) in cell culture media. **(a)** pH of the NPC culture medium was approximately neutral (∼pH 7.5) both before and after incubation in the CO₂ incubator (Methods). **(b)(c)** Distributions of zeta potential and size (bubble diameter) for NB1 and NB6. See Table 1 for details of NB1 and NB6. Each line in each figure represents NBs measured at a different time interval using the NanoSight instrument (Malvern Instruments) (Methods).

**Table 1.** Summary of the physical properties of the nanobubble (NB)-containing cell culture media.

| NB<br>medium | Zeta potential<br>(mV) | Concentration<br>($\times 10^8$ particles/mL) | Medium<br>preservation |
| --- | --- | --- | --- |
| NB1<sup>a</sup> | +12.2± 0.8 | 1.9±2.4 | 1 month |
| NB2<sup>a</sup> | +25.9± 0.7 | 4.4± 0.3 | 1 week |
| NB3<sup>b</sup> | +26.9± 2.1 | 1.0±0.3 | 1 week |
| NB4<sup>b</sup> | +37.1 ± 5.0 | 1.6±2.2 | 1 week |
| NB5<sup>a</sup> | -5.3 ± 0.3 | 7.7±1.6 | 1 week |
| NB6<sup>a</sup> | -14.8± 0.1 | 3.6±0.2 | 1 month |

a: cell culture medium for human iPSC-derived NPC
b: cell culture medium for human iPSC-derived neurons

We prepared six types of NB-containing media using cell culture media for iPSC-derived cells (Table 1) (Methods). In this study, approximately 20 mL of nanobubble-containing culture medium was generated in a single preparation. At present, the physical properties of NBs generated in the culture media using the patented charge activation plate contact method (see patent numbers in the Methods section), specifically their concentration and zeta potential, were not fully controlled and showed some variability (Table 1). For NB1 and NB6 media, Figures 1b and 1c show the distributions of zeta potential and particle size, measured by using the NanoSight system (Malvern Instruments) (Methods). We found that NB1 and NB6 had similar physical properties and were suitable for comparison. In the NanoSight system, particle size is determined from the mean squared displacement of scattered light signals using the Stokes–Einstein equation. Zeta potential is measured by applying an electric field between two platinum electrodes, calculating electrophoretic mobility from particle velocity (corrected for electro-osmotic flow), and converting it using the Smoluchowski approximation. The NanoSight has been used in NB research^5^. The nanoparticle tracking analysis (NTA) of the NanoSight has been compared with dynamic light scattering (DLS), and its accuracy has been evaluated.^15–17^ Although the distributions of zeta potential were broad (Figs. 1b and 1c, top), it was evident that NB1 medium primarily contained positively charged NBs, whereas NB6 medium mainly contained negatively charged NBs. According to the size distributions, most of the particles were in the submicron range (Fig. 1b and 1c, bottom), confirming that they qualified as NBs in accordance with the international standard ISO/TC281.

Apart from NTA analysis, the dynamic properties of NBs were evaluated by dark-field microscopy, which revealed their Brownian motion in the culture medium. Their diffusional behavior for 10 seconds was consistent with the estimation from the Einstein relation for a submicron-scaled particle (Fig. S1 in Supplementary Information).

### Physicochemical properties of NB-free cell culture medium

The experimental results using NanoSight for the medium without NBs are shown in Fig. 2. The zeta potential is different from those in Fig. 1, indicating that the results in Fig. 1 arise from the presence of NBs. Because the culture medium contains biomolecules, some components were detected as particles; however, the zeta potential was, on average, close to zero (+2.5±1.4 mV). Additionally, when the culture medium was frozen to disrupt NBs and subsequently reanalyzed by NTA, a mean zeta potential was −2.1±0.5 mV. The fact that the mean zeta potential before and after freeze-thaw treatment remains the same within experimental error indicates that charged NBs were scarcely present in the original culture medium. Regarding the particle concentration obtained from NTA by using the NanoSight system, Δparticle concentration before and after freeze-thaw treatment was −170 %. It indicates that the detected particle number was at the noise level where noise arose from the setting of the brightness threshold for particle detection, and that the number of particles may increase after freeze–thaw treatment due to fragmentation of components in the culture medium.

**Fig. 2.**
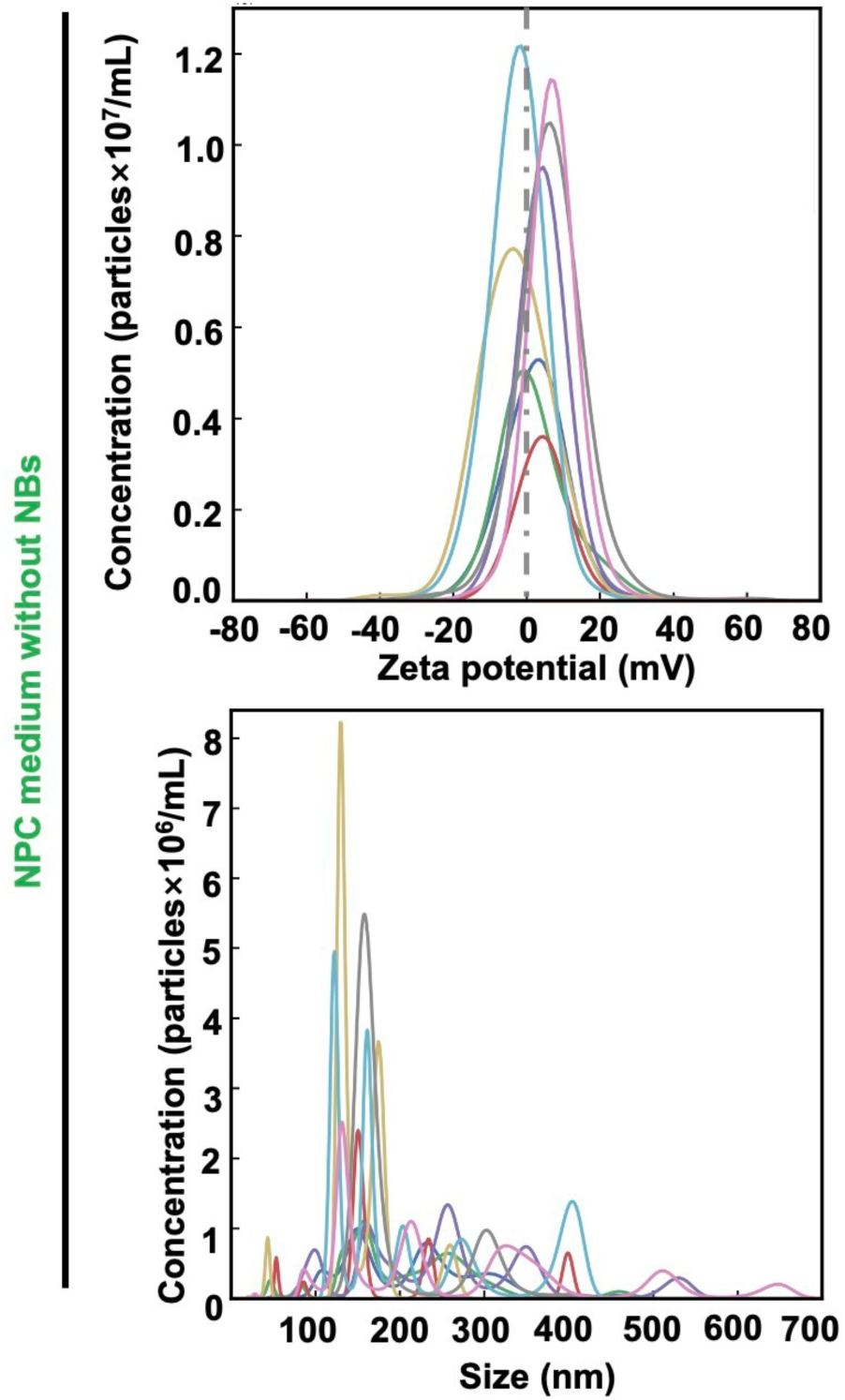
Physical properties of nanobubbles (NBs)-free cell culture media. The NB-free iPSC-derived NPC culture medium was examined by using the NanoSight instrument (Malvern Instruments) (Methods) for comparison. Each line in each figure represents NBs measured at a different time interval using the NanoSight instrument.

### Culture and observation of neural progenitor cells (NPCs)

Based on the timeline of the culture (Fig. 3a), on the fifth day of starting the culture of the human iPSC-derived NPCs (Methods), the expression of NPC-specific transcription factors SOX2 and PAX6^18^ was confirmed using the immunocytochemistry (ICC) method (Methods) (Fig. 3b).^19^ We defined the day as “Day 0” and performed subsequent observations using phase-contrast microscopy over the following three days (Fig. 3c). In Fig. 3c, because the NPCs underwent cell division, we observed a gradual increase in cell density in the absence of NBs in the medium.

**Fig. 3.**
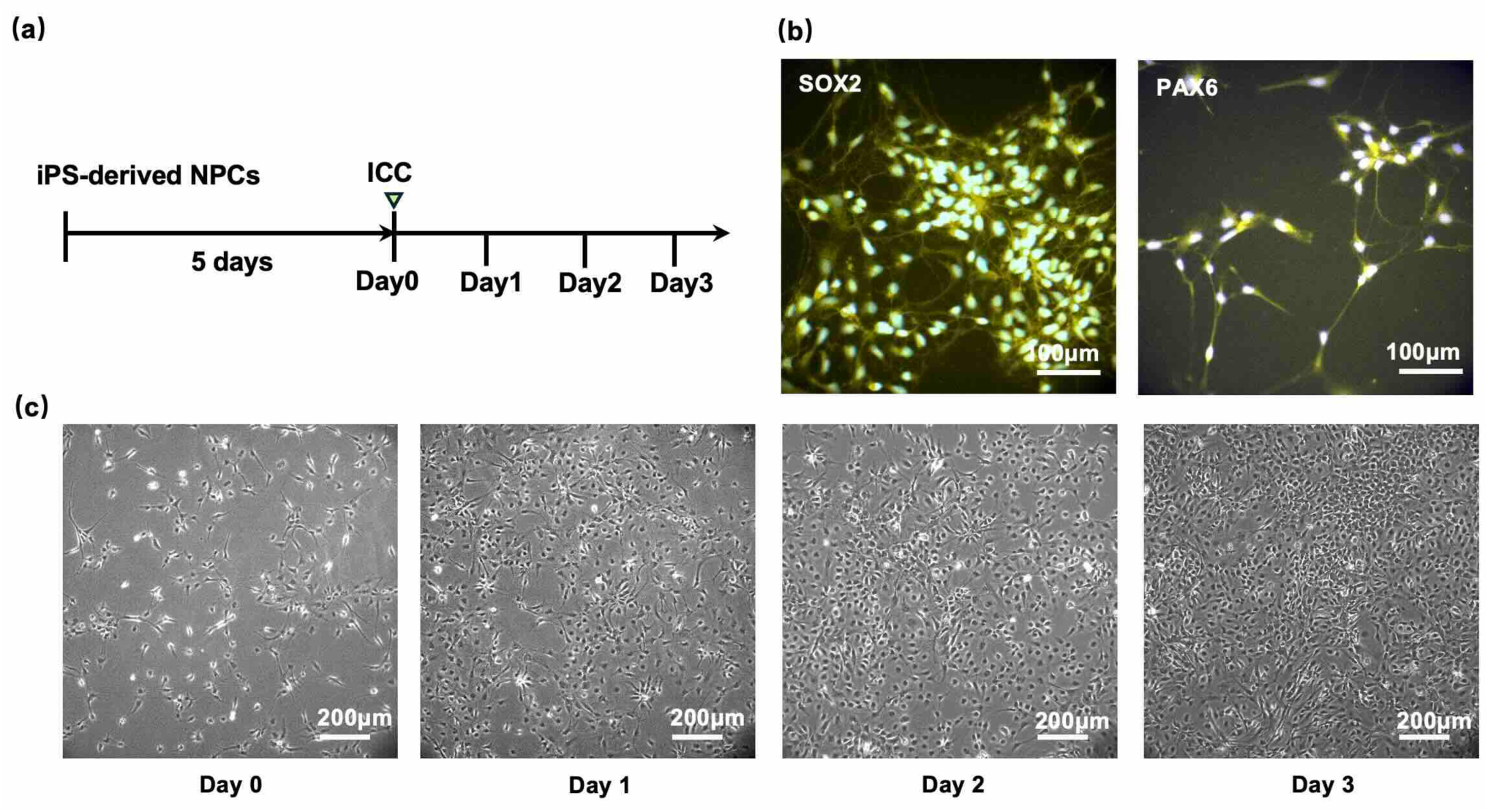
Culture of human iPSC-derived NPCs without nanobubbles (NBs). **(a)** Timeline of cell culture, ICC, and fluorescence observation. **(b)** Immunocytochemistry (ICC) fluorescence images of human iPSC-derived NPCs stained for the markers SOX2 and PAX6 using a secondary antibody conjugated with Cy3 (Methods). Scale bars: 100 µm. **(c)** Time-lapse phase-contrast images of NPCs over three consecutive days of culture. Day 1, Day 2, and Day 3 correspond to 24, 48, and 72 hours after Day 0, respectively. Live cells appear dark because they adhered to the glass surfaces. The cell density increased over time. Scale bars: 200 µm.

The live NPCs appeared phase dark because they adhered to the coverslip exhibiting typical morphology. In contrast, dead cells appeared phase bright because they shrank in volume, rounded up, and fell out of the primary focal plane (phase contrast images in Fig. 4a). Using a cell viability assay we confirmed the correlation between cell morphology and viability (Fig. 4a). Live cells adhered to the glass surface and appeared dark; they stained with Calcein AM but not with EthD-III, while dead cells lost adhesion, appeared phase bright, and stained with EthD-III but not with Calcein AM. Here, the nuclei of dead cells exhibited strong fluorescence and showed clearly different fluorescence intensities (FIs) compared to live cells (Fig. 4b; *p*=7.7× 10^−68^, two-tailed Student’s T-test). The increased fluorescence in dead cells is primarily due to a sequence of events: enhanced membrane permeability, influx of nucleic acid-binding dyes, fluorescence enhancement upon binding, and cessation of dye efflux. Therefore, cells with an FI below 100 (a.u.) were counted as live cells later when quantifying the number of viable cells. Because cells were occasionally overlapping, the detection of live cells using our particle-detection software (described later) was better suited with Hoechst-stained images that detected the nuclei than with those stained with Calcein AM, as Calcein AM stains the entire cytoplasm. For this reason, live-cell counting was performed using Hoechst staining in this study. The correlation between Hoechst imaging and EthD-III imaging, which also detected nuclei, was further examined in the Supplementary Information (Fig. S2), where their consistency was confirmed. In addition, previous studies have reported that Hoechst imaging brightness correlates with cellular damage. In a previous study, using H_2_O_2_ treatment, nuclear condensation was monitored with DNA-chelating dyes such as Hoechst, and apoptosis was discussed based on the increased fluorescence intensity of Hoechst staining.^20^ Another study also examined Hoechst imaging in combination with DiOC6 and YO-PRO-3 imaging, demonstrating that Hoechst-bright cells exhibit nuclear condensation and loss of mitochondrial function.^21^

**Fig. 4.**
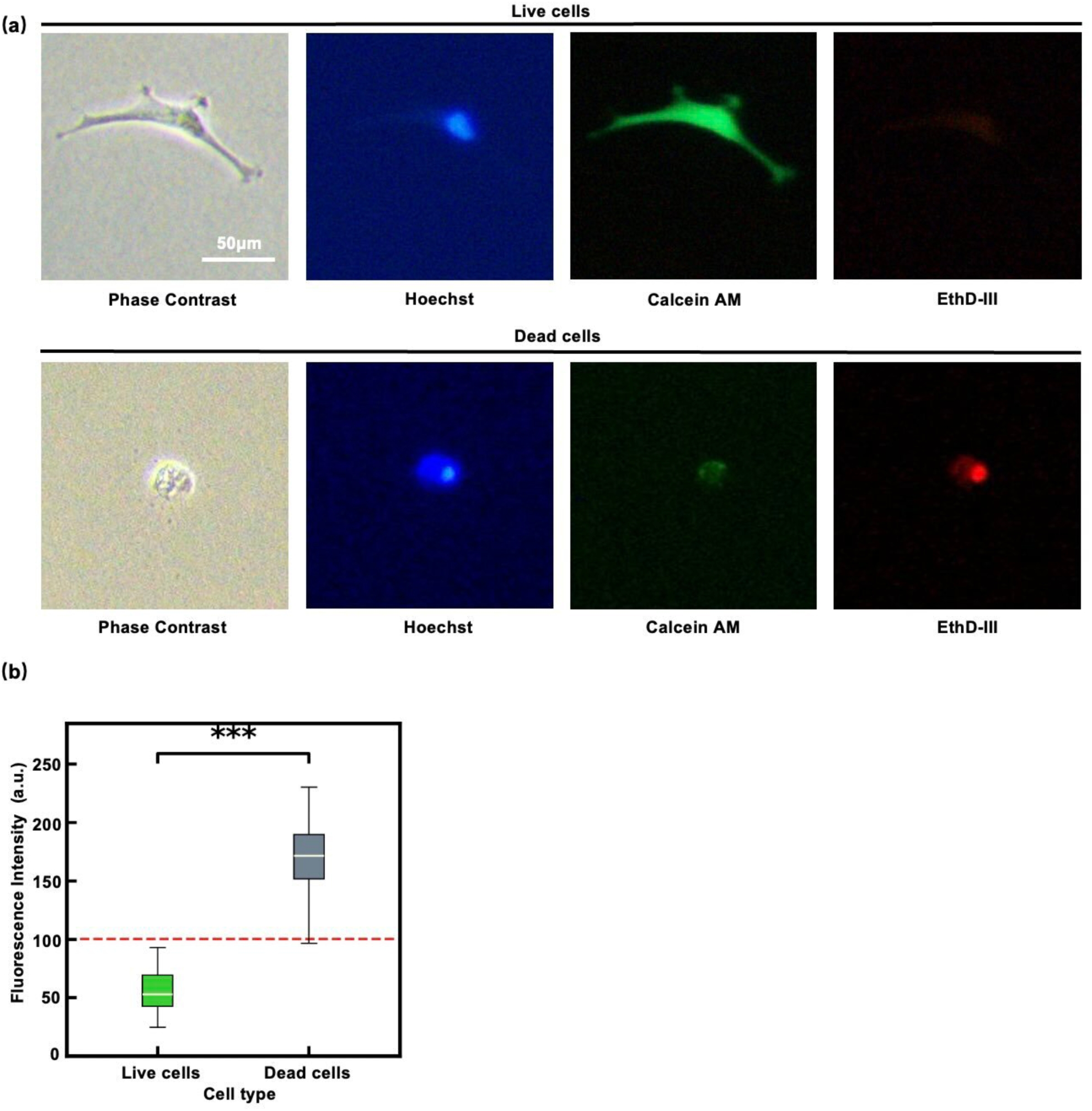
Live/Dead cell assay of NPCs. **(a)** Representative images of live/dead NPCs stained with Hoechst (blue), Calcine AM (green), and EthD-III (red). Here, Hoechst stains cell nuclei, Calcein AM labels live cells, and EthD-III stains dead cells. The shrunken cells observed under phase-contrast microscopy, showing low fluorescence intensity (FI) for Calcine AM and high FI for EthD-III, were dead, whereas live cells showed the opposite pattern. Scale bar: 50 µm. **(b)** Comparison of fluorescence intensities of live and dead cells (*p*=7.7× 10^−68^, two-tailed Student’s T-test). The dead cells exhibit strong fluorescence, and they can be distinguished from the live cells. Statistical significance is indicated as follows: \**p* < 0.05, \*\**p* < 0.01, and \*\*\**p* < 0.001. The error-bars represent SD.

### Addition of NB-containing NPC culture media

On the fifth day of starting the culture, after the expression of the transcription factors was confirmed (Fig. 3b), the culture medium was replaced with the NB-containing media. Note that the NB-containing media used here (NB1 and NB6 media in Table 1) were prepared approximately one month (43 days) before the experiment. NB1 and NB6 had similar physicochemical properties, as shown in Fig. 1 and Table 1, and were used as comparative samples. In a later section, the effects of the culture media (NB2 and NB5) immediately after NB generation were also investigated.

The timeline of the following experiments is shown in Fig. 5a. The observation period after NB addition (3 days) was determined with reference to a previous study on fibroblasts using NBs.^22^ In Fig. 5b, NPCs were observed using phase-contrast microscopy from Day 1 to Day 3. Furthermore, the nuclei of NPCs were stained with Hoechst^23^ (blue) to facilitate later detection of the cells and enable cell counting. In the NB1 medium containing positively charged NBs, the cells shrank and rounded up due to loss of adhesion to the glass surface as the days passed. The NB1 medium containing positively charged NBs rapidly reduced the number of live NPCs, whereas the NB6 medium with negatively charged NBs caused a gradual decrease in the live cell numbers (cell counts are provided in the next section). Note that the results here are entirely different from those in Fig. 3c, where continuous cell growth was observed in the absence of NBs. These findings indicate that NBs generated nearly one month prior to use still had a significant effect on the cells. Such a cytotoxic effect has also been reported in fibroblasts induced by hydroxyl radicals.^14,22^ Moreover, there was a clear difference between positively and negatively charged NBs, an issue that previous studies did not address because they were unable to generate positively charged NBs. The clear difference may be related to the fact that positively charged NBs generate more hydroxyl radicals, or to the negative charge of the cell membrane (approximately −100 mV to −10 mV),^24^ which could lead to electric repulsion of negatively charged NBs.^2,25^

**Fig. 5.**
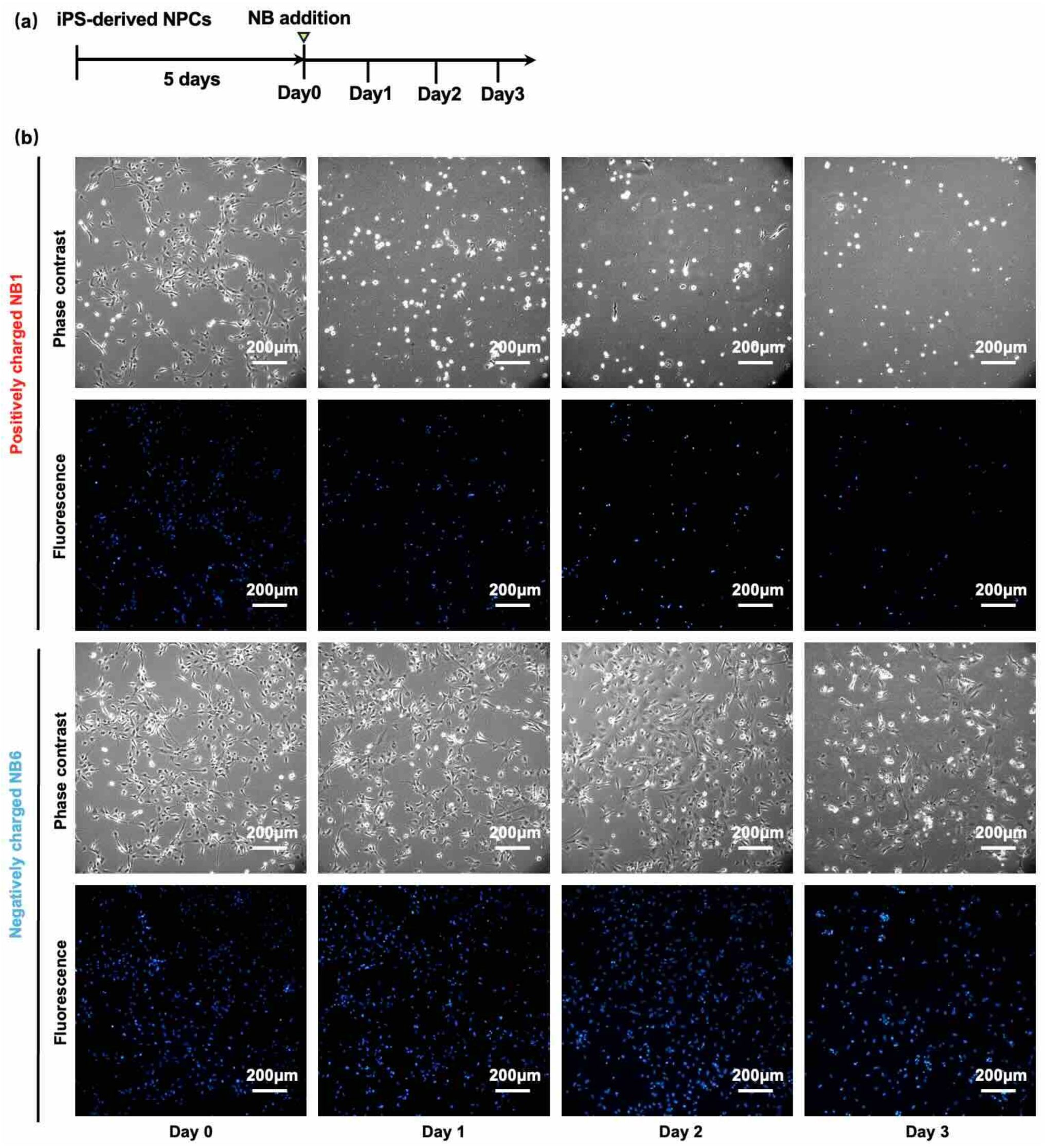
Culture of human iPSC-derived NPCs with nanobubbles (NBs). **(a)** Timeline of NB-medium addition experiments. **(b)** Phase-contrast and fluorescence images, where the nuclei of cells were stained with Hoechst, showing the culture of NPCs over three days in NB1 (top) and NB6 (bottom) media (Table 1). Scale bars: 200 µm.

### Counting the number of live NPCs

To quantify the number of live NPCs, their nuclei were stained with Hoechst^23^, making them clearly visible and easier to detect as particles in the image analysis (Fig. 6a). Although Calcein AM is appropriate for identifying live cells (Fig. 4a), cell detection becomes difficult when cells overlap. Therefore, we quantified the number of live cells by detecting nuclei using Hoechst staining (Fig. 6b, left) using our custom particle detection software (Figs. 6a, bottom and 6b, right). The distinction between live and dead cells was made using the criteria shown in Fig. 4b, with the correlation to the Live/Dead cell assay using Calcein AM/EthD-III examined in the Supplementary Information (Fig. S2). In this step, the nuclei of the NPCs were identified using the software by scanning the entire image with a nucleus-sized ROI (region of interest) (Figs. S3-S5 of the Supplementary Information). In the previous studies,^14,22^ cells were counted manually or by using ImageJ (NIH), a general-purpose image analysis program, whereas we developed software based on the overlapping ROI scanning method (Figs. S3-S5 in the Supplementary Information).

**Fig. 6.**
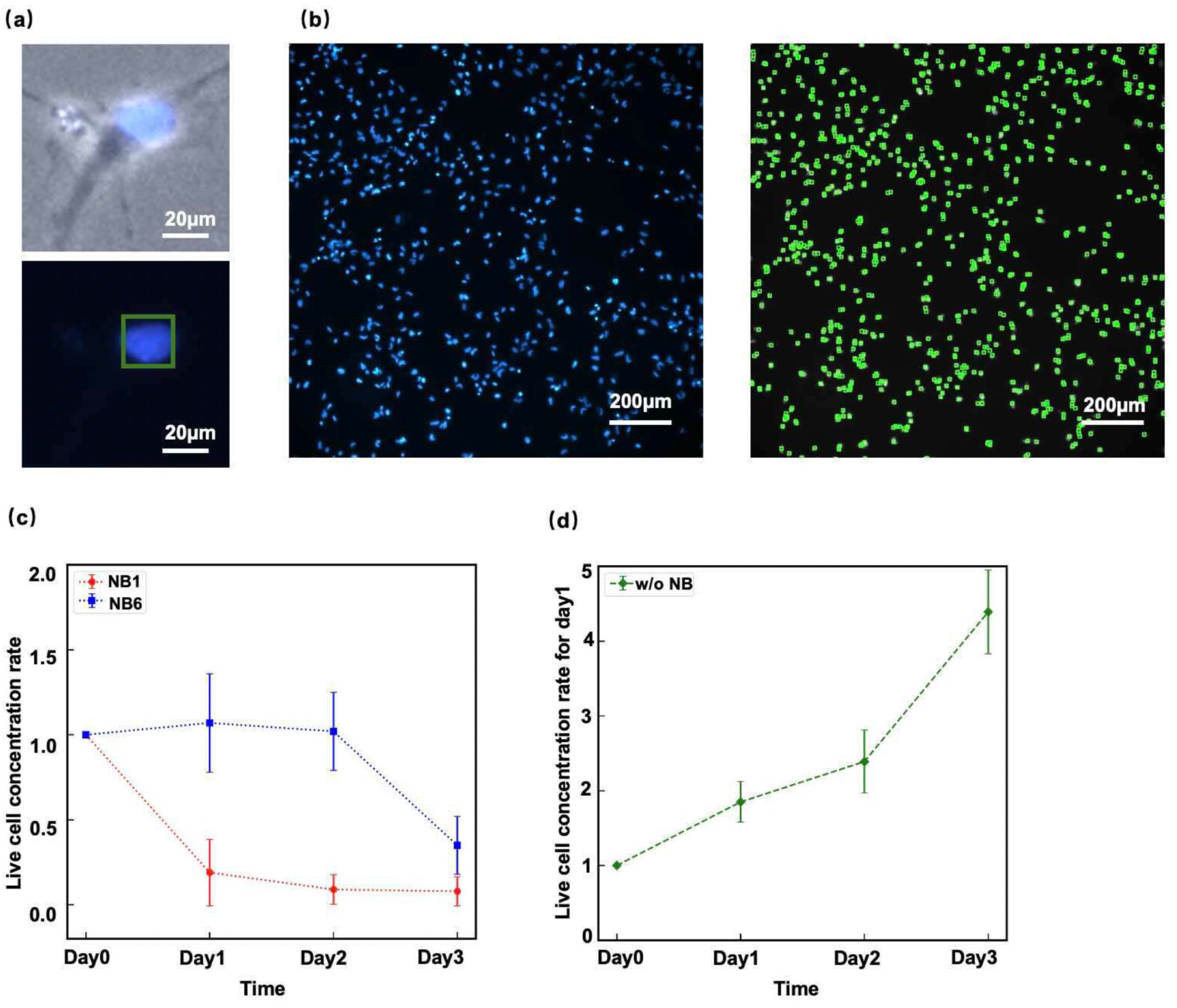
Quantification of nanobubble (NB) effects on NPCs by live/dead cell counts. **(a)** Example of cell counting using our custom particle detection software (Figs. S3-S5 of the Supplementary Information). Cell nuclei were stained with Hoechst, and cells were detected based on their fluorescence intensities (FI). Scale bars: 20 µm. **(b)** The number of NPCs in a single fluorescence image, covering an area of 1 mm^2^, was automatically calculated by the software. For each day and nanobubble (NB) condition, 10 images were randomly taken to observe the cells plated in a well of a 6-well plate (area: ∼10 cm²). Scale bars: 200 µm. **(c)** Live cell concentration rates (Eq. 1) for *N* = 0,1,2,3 in the cases of the positively charged NB1 (red) and negatively charged NB6 (blue) media (Table 1), respectively. The error-bars represent SD. **(d)** Live cell concentration ratio (Eq. 1) for *N* = 0,1,2,3 in the absence of NBs (Fig. 3c). NPCs proliferated in the absence of NBs.

A single microscopic image covers an area of 1.77 mm^2^, and 10 images were randomly taken to observe the cells plated on the dish, whose area was approximately 10 cm^2^. Note that detailed protocols for the cell experiments are described in Fig. S6 of the Supplementary Information), which also indicates that a total of seven independent experiments were performed in this study. Then, the live cell ratio (LCR) as a function of the *N*-th day of culture was defined as:

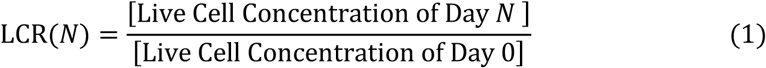

Normalization to Day 0 values was performed to minimize variability in cell density. Here, the initial cell density is approximately 10⁵ cells/well. The calculated values of LCR(*N*) for *N* = 0, 1, 2, 3 (*N* denotes the number of days passed) are shown in Fig. 6c for the positively charged NB1 medium (red) and negatively charged NB6 medium (blue), respectively. Note that by definition, LCR(0) = 1. When NPCs were cultured in the medium without NBs (Fig. 3c), LCR(*N*) increased (Fig. 6d), whereas in the media containing NBs, LCR(*N*) decreased over time (Fig. 6c). The decrease in LCR(*N*) in the presence of NBs was more pronounced with positively charged NBs (NB1) (red line in Fig. 6c) than with negatively charged NBs (NB6) (blue line in Fig. 6c).

### Effect of NB storage time on cell viability

In Table 1, NB2 and NB5 media are those in which NBs were generated immediately before use, and at most about a week had passed during the experiments. Figures 7a and 7b show the zeta potentials and size distributions of the NB2 and NB5 media, respectively. These distributions were measured using the NanoSight (Methods). Similar to the media prepared one month after NB generation (NB1 and NB6 media), both positively and negatively charged NBs exhibited high zeta potential values (several tens of mV).

**Fig. 7.**
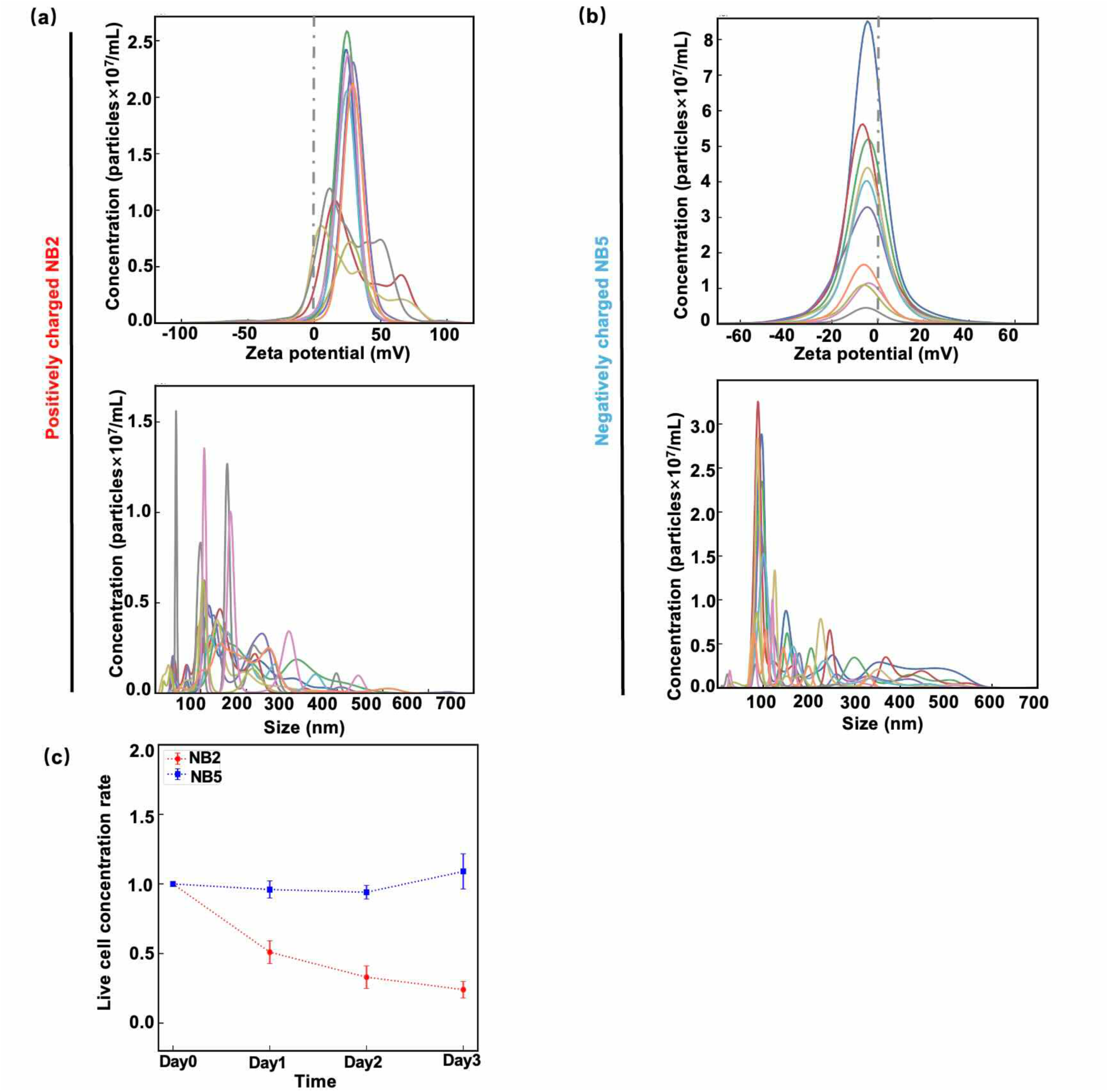
Nanobubble (NB) addition experiments on NPCs using NB2 and NB5 media. **(a)(b)** Distributions of zeta potential and size for NB2 and NB5. See Table 1 for the details of NB2 and NB5. Each line represents NBs measured in a different time interval using the NanoSight instrument (Malvern Instruments) (Methods). **(c)** Live cell concentration ratios (Eq. 1) for *N* = 0,1,2,3 in the cases of the positively charged NB2 (red) and negatively charged NB5 (blue) media (Table 1), respectively. The error-bars represent SD.

We also conducted cell viability experiments using NB2 and NB5 media. Figures 7c shows the LCR(*N*) defined by Eq. 1 for the cases of NB2 and NB5 media. The behaviors of LCR(*N*) for the cases of the NB2 and NB5 media were similar to those for NB1 and NB6 media (Fig. 6c). Note that the superposition of these graphs (graphs in Figs. 6c and 7c) is provided in Fig. S7 of the Supplementary Information, demonstrating their similarity. The results suggest that NB stability and their cellular effects remain largely unchanged between one week and one month after NB generation in the media, as confirmed by the cell viability experiments.

### The case for human iPSC-derived neurons

For the NB addition experiment targeting neurons, the iPSC-derived NPCs were differentiated into forebrain neurons, and the culture process along with the experimental timeline is presented in Fig. 8a. The NPCs were differentiated into neurons as previously described (Methods)^19^ and after two weeks, neuronal maturation and neurite outgrowth were confirmed by phase-contrast microscopy (Fig. 8b). Additionally, the expression of neuron-specific proteins such as βIII-tubulin^26^ and Tau^27^ was confirmed by ICC (Fig. 8c) (Methods) indicating that the NPCs successfully differentiated into neurons. Then, Day 0 was set as the second week after the differentiation, and the medium was replaced with the NB3 medium, whose zeta potential was similar to that of NB2 medium for NPCs (Table 1). Here, we also succeeded in generating NBs not only in the medium for NPCs but also in the maturation medium for iPSC-derived neurons. The results of live cell counting are shown in Fig. 8d, where LCR(*N*) (Eq. 1) for *N* = 0, 1, 2, 3 are plotted. A decrease in live cell number was observed due to the influence of the positively charged NB-containing medium. However, the effect of positively charged NBs was not as significant as that observed in NPCs. To further support these findings, we generated NB4 medium (Table 1). Even though NB4 medium exhibited a higher zeta potential than NB3 medium, its effect was not significant (Fig. 8e).

**Fig. 8.**
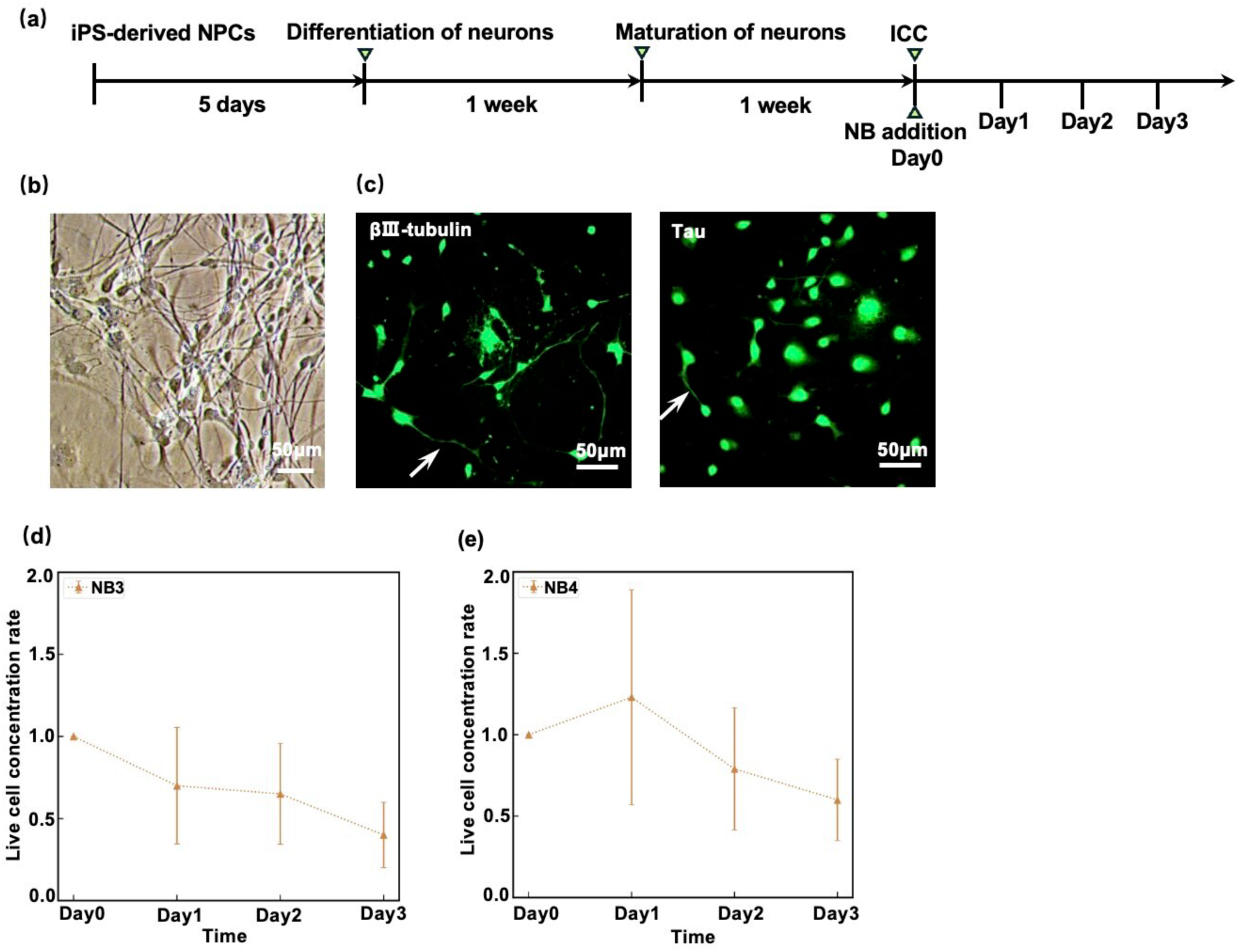
Culture of human iPSC-derived forebrain neurons with nanobubbles (NBs). **(a)** Timeline of NB-medium addition experiments for the neurons. **(b)** Phase-contrast images of human iPSC-derived neurons. Two weeks after the differentiation process (Methods), neurites had extended and the morphology had become neuron-like. Scale bar: 50 µm. **(c)** Immunocytochemistry (ICC) fluorescence images of the neurons stained with the markers βIII-tubulin and Tau, using secondary antibodies conjugated to Alexa Fluor 488 (Methods). White arrows indicate axons and dendrites. Scale bars: 50 µm. The arrows indicate processes characteristic of neurons. **(d, e)** Live cell concentration ratios (Eq. 1) for *N* = 0,1,2,3 in the cases of NB3 and NB4 media, respectively (Table 1). The error-bars represent SD.

### Summary and discussion

In this study, we demonstrate that both positively and negatively charged nanobubbles (NBs) can be stably generated in neutral solvents around pH 7, particularly in media used for the culture of iPSC-derived cells, and we provide supporting evidence for their stability (Figs. 1 and 7). All results regarding the application of NB-containing media (Table 1) to the cells are summarized in Fig.9. Because our NBs remained stable for up to one month, we next attempted to generate them in cell culture media for iPSC-derived neurons, which typically require several weeks to approximately one month for differentiation. Importantly, we successfully generated not only negatively charged NBs but also positively charged NBs in the neuronal culture medium, which had previously been difficult to maintain in a stable state^14^. We found that the positively charged NBs induced stronger cytotoxic effects in the iPSC-derived cells (Fig. 9).

**Fig. 9.**
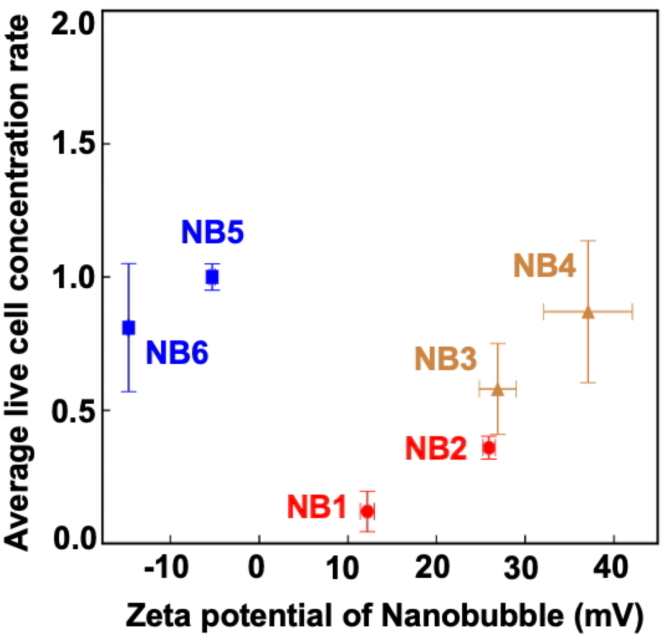
Summary of effect of nanobubbles (NBs) on human iPSC-derived NPCs and forebrain neurons. NB1 to NB4 represent positively charged NB-containing cell culture media (red and orange), whereas NB5 and NB6 are negatively charged (blue). NB3 and NB4 were applied to the neurons, while NB1, NB2, NB5, and NB6 were applied to the NPCs. The *y*-axis represents the 3-day average of the live cell concentration ratios (Eq. 1). The similar values obtained for NB1 and NB2, NB3 and NB4, and NB5 and NB6 demonstrate the reproducibility of the results. The error-bars were calculated based on the standard error propagation formula (Eq. 4).

Regarding the finding that positively charged NBs had a greater effect than negatively charged NBs for iPSC-derived cells (Figs. 6c and 7c), one possible reason is that positively charged NBs may generate stronger radical production than negatively charged NBs. This point needs to be examined in more detail in future studies by analyzing the radicals generated. Another possible reason is that the cell membrane is negatively charged (from −100 mV to −10 mV).^24^ Because negatively charged particles may be less likely to approach cells, it is considered that uptake into cells via endocytosis and phagocytosis is considered to be limited.^2,25^ This point has also been noted in previous studies on the application of NBs to *Helicobacter pylori* cells and fibroblasts.^2,14^ Although not NBs, similar findings have been reported for silica nanoparticles. In particular, the study have shown that 300-nm nanoparticles with a zeta potential of +40 mV exhibit greater cellular uptake than those with −40 mV.^28^ Fundamentally, the negatively charged cell membrane functions as a natural barrier to the entry of polyanionic nucleic acids, such as negatively charged DNA molecules, which are recognized as foreign molecules^29^.

In the case of neurons (Fig. 8d), the NB3 medium did not reduce LCR(*N*) (Eq. 1) as much as in NPCs (Fig. 6c), despite having similar physical properties, such as zeta potential and concentration, to those of the NB2 medium (Table 1). One possible explanation is that terminally differentiated cells, such as neurons, have less active endocytosis than proliferating cells (including NPCs), which may lead to reduced uptake of NBs and consequently a smaller impact.^30^ If NBs could be used to selectively eliminate unwanted proliferating cells, such as glial cells, while minimizing their impact on neurons in iPSC-derived neuron cultures, this approach could improve the quality of neuron cultures and be beneficial for regenerative medicine applications involving iPSC-derived neurons. It is important to elucidate, from a molecular biological perspective, the differences in cellular sensitivity to NBs across cell types.

## Materials and Methods

### Nanobubble (NB)-supplemented medium

The positively or negatively charged NB-supplemented culture media were prepared using the original charged NB production method developed by the co-author, Ohdaira (Ohdaira Laboratory Co., Ltd., Saitama, Japan) (Japanese Patent Nos. 7637707 and 7227694; US Patent No. 12329152; US Application No. 16/770528; Chinese Patent No. ZL201880088629.2; Korean Patent Nos. 10-2667835 and 10-2657332; European Patent No. 3721886). In this paper, the characteristics of the charged NBs were described based on data obtained from the NanoSight NS500 system (Malvern Panalytical Ltd., UK) (Figs. 1b, 1c, 2, 7a, and 7b). Using the developed NB production method, termed the “charge activation plate contact method,” positively or negatively charged NBs were generated directly in cell culture media without introducing any externally added charging agents or applying electric currents. Two key characteristics of the NB production method are: (1) NBs can be directly charged within a solvent (cell culture medium) without any intervening substances between the solvent and the charged activation plate used as the NB generator, and (2) the desired charge polarity is maintained immediately upon NB generation. In this study, NBs were specifically generated in STEMdiff™ Neural Progenitor Medium (STEMCELL Technologies, Cat#05833; corresponding to NB1, NB2, NB5, and NB6 in Table 1) and STEMdiff™ Neuron Maturation Medium (STEMCELL Technologies, Cat#08605; corresponding to NB3 and NB4 in Table 1). At present, 20 mL of culture medium can be prepared using the activation plate, which was the amount used for a single independent culture. In addition, the NB concentration and zeta potential showed variability and were not yet fully controlled (Table 1).

### Nanoparticle tracking analysis (NanoSight NS500)

Zeta potentials and sizes of NBs were measured using the NanoSight NS500 system (Malvern Panalytical Ltd., UK), equipped with the Z-NTA (Zeta–Nanoparticle Tracking Analysis) module and controlled via the Z-NTA analysis software (NTA 3.2). 500 µL of NB-containing medium was loaded into the sample chamber using a syringe connected to the sample port. After flushing the system with clean diluent, the sample was introduced into the flow cell, and measurements were conducted at room temperature. The NanoSight NS500 combines a laser-based dark-field optical system with Z-NTA to determine particle size and zeta potential on a particle-by-particle basis. Scattered light from Brownian particles is recorded by a camera, and particle size is calculated from the mean squared displacement via the Stokes–Einstein equation. Zeta potential was obtained from electrophoretic mobility, measured under an applied electric field, with electro-osmotic flow corrected by multi-depth velocity profiling, using the Smoluchowski approximation.

### pH measurement

The pH of STEMdiff™ Neural Progenitor Medium (STEMCELL Technologies, Cat#05833) was measured using a LAQUAact pH Meter (HORIBA Scientific, D-71) three times at room temperature under two conditions: one was equilibrated overnight in a 37 °C, 5% CO₂ incubator, and the other was equilibrated outside the CO₂ incubator.

### Dark field microscopy

Dark-field microscopic observation of NBs in the cell culture medium (Fig.S1 of the Supplementary Information) was performed using an inverted microscope (ECLIPSE Ti2-E, Nikon, Tokyo, Japan) equipped with a dark field condenser (Cat#MBL12010, Nikon Corporation, Tokyo, Japan) and a 100× oil immersion objective (CFI Plan Fluor 100×, Cat# MRH01902, Nikon Corporation, Tokyo, Japan). Images were recorded at 33 frames per second with pixel size of 1.3 µm by a CMOS camera (ORCA-Flash 4.0 V2, Hamamatsu Photonics). Here, the NB-containing medium was loaded into a prepared chamber consisting of two cover glasses (24 × 24 mm and 24 × 32 mm; Matsunami Glass Ind., Ltd., Cat# C022430 and C022432), separated by two spacers of ∼50 µm thickness, for observation.

### Preparation for human iPSC-derived NPCs

The human iPSC-derived Neural Progenitor Cells (NPCs) line was obtained from STEMCELL Technologies (Cat#200-0620). The human iPS-derived NPCs were cultured and expanded based on the established protocols^19^ with minor modifications. The surface of a 6-well plate (Thermo Fisher Scientific, Cat#140675) was coated with Corning Matrigel Matrix (Corning, Cat# 3038002), diluted in DMEM/F12(Thermo Fisher Scientific, Waltham, MA, USA, Cat# 21041025) supplemented with 15 mM HEPES (Gibco, Thermo Fisher Scientific, Waltham, MA, USA, Cat# 15630106). Cryopreserved NPCs were thawed at 37 °C, centrifuged (300 × g, 5 min, RT), and the supernatant discarded. The cell pellet was resuspended in STEMdiff™ Neural Progenitor Medium (STEMCELL Technologies, Cat#05833). The cells were then plated onto the Matrigel-coated 6-well plate at a density of 6.0 × 10⁵ cells/well, counted using the EVE™ Automatic Cell Counter (NanoEnTek Inc., NESMU-EVE-001E), and incubated at 37°C in a humidified 5% CO₂ incubator. The medium was replaced every two days (1 mL/well). NPCs were passaged once they reached approximately 70–80% confluency, typically 5–7 days after seeding, using 1 mL/well Accutase (STEMCELL Technologies, Cat#07920) at 37 °C for 5 min for detachment. After neutralization of Accutase with 2 mL/well DMEM/F12 and gentle mechanical dissociation, cell suspensions were centrifuged under the same conditions, resuspended in the fresh medium, and then seeded onto newly coated plates at an appropriate split ratio for expansion. All experiments in this paper were performed using NPCs at passages below 4. One vial of NPCs purchased from STEMCELL Technologies was expanded to generate multiple stocks of cells (vials) at passage 4 or lower. Seven independent experiments were performed using one vial of expanded cells at passage 4 or lower for each of the seven culture media: NB-free medium and media containing NB1 through NB6 (Table 1). A schematic of the experimental workflow is provided in Fig. S6 of the Supplementary Information.

### Preparation for human iPSC-derived forebrain neurons

iPSC-derived forebrain neurons used in this paper were differentiated from the human iPSC-derived NPCs using STEMdiff™ Forebrain Neuron Differentiation Medium (STEMCELL Technologies, Cat# 08600), based on the established protocols^19^ with minor modifications. After exchanging the differentiation medium for 2mL/well, the cells were fed every 2-3 days with the differentiation medium and incubated for 5 days. For neuronal maturation, differentiated neurons were again detached using Accutase (STEMCELL Technologies, Cat#07920), washed, and resuspended in STEMdiff™ Neuron Maturation Medium (STEMCELL Technologies, Cat# 08605). Subsequently, the neurons were transferred onto plates coated with PLO (50 µg/mL; Sigma-Aldrich, Cat# P3655) and Laminin (5 µg/mL; Sigma-Aldrich, Cat# L2020) and seeded at a final density of 4.0 × 10⁵ cells/mL, counted using the EVE™ Automatic Cell Counter (NanoEnTek Inc., NESMU-EVE-001E). Cells were maintained at 37 °C in a humidified 5% CO₂ incubator. Half of the maturation medium was changed every 2–3 days. The neurons were used for observation after differentiation for around 2 weeks.

### Immunocytochemistry (ICC)

The ICC experiments in this study were performed based on the established protocols^19^. For the ICC experiments, cells were cultured on coverslips (18 × 18 mm, IWAKI, Cat# 62512; AGC Techno Glass, Japan) after the first passage for NPCs and after maturation stage for neurons. Cells were fixed with 4% paraformaldehyde (PFA; FUJIFILM Wako, Cat# 163-20145) at 37 °C for 15 min. After fixation, the cells were rinsed twice with Dulbecco’s phosphate-buffered saline (DPBS; Gibco, Cat# 14080055) for 1 min each. Permeabilization was carried out in 0.25% Triton (Nacalai Tesque, Cat# 3550115) in DPBS for 5 min at room temperature, followed by three 5-min times washes in DPBS. Non-specific binding of antibodies to the glass surface was blocked in 0.5% fish-skin gelatin in PBS at 37 °C in a humidified 5% CO₂ incubator for 1 h. Primary antibodies (PAX6 and SOX2 for NPCs; Tau and βIII-tubulin for neurons) were diluted in the blocking solution at the following concentrations and applied to cells overnight at 4 °C: mouse anti-PAX6 (1:500, Developmental Studies Hybridoma Bank, Cat# PRB-278P), mouse anti-SOX2 (1:200, DSHB, Cat# PCRP-SOX2-1B3), mouse anti-Tau (1:500, DSHB, AB_528487) and mouse anti-βIII-tubulin (1:500, DSHB, AB_528497). Secondary antibodies were diluted 1:500 in the blocking buffer and incubated with the cells for 1 h at 37 °C in the dark. The following secondary antibodies were used: goat anti-mouse IgG (H+L), Cy3-conjugated (1:500, Jackson ImmunoResearch, Cat# 115-165-146, Ex/Em: 545/605 nm) for SOX2 and PAX6; and goat anti-mouse IgG (H+L), Alexa Fluor 488-conjugated (1:500, Thermo Fisher Scientific, Cat# A11001, Ex/Em: 470/525 nm) for the neuron-specific markers Tau and βIII-tubulin. Coverslips were washed in DPBS. Coverslips were mounted onto glass slides (25 × 75 mm, 1.0–1.2 mm thick, ASLAB) using ProLong™ Gold Antifade Mountant with DAPI (Thermo Fisher Scientific, Cat# S36920), and then stored at 4°C overnight before imaging.

### Live and dead cell viability assays

Cell viability was assessed using a Live/Dead assay kit (Biotium Cat # 30002, Fremont, CA, USA). Cells were incubated with 4 µM ethidium homodimer III (EthD-III, Ex/Em: 279, 532/625 nm) and 2 µM calcein AM (Ex/Em: 494/517 nm), which respectively label dead and live cells, for 5 min in the CO_2_ incubator. Then cells were washed once with 1 mL Hank’s balanced salt solution (Thermo Fisher Scientific, Cat# 14025-092). Cell images were acquired by fluorescence microscopy (see the “Fluorescence observation” section below).

### Nucleus observation

Hoechst 34580 (Thermo Fisher Scientific, Cat# S0486, Ex/Em: 360/461 nm) was used for nuclear staining at a final concentration of 10 µg/mL. Hoechst 34580 was added to the plate and the cells were incubated for 5 min in the incubator. After incubation, the staining medium was removed, and the cells were washed once with 1 mL of DMEM/F12. Subsequently, 1 mL of fresh DMEM/F12 was added to each well of a 6-well plate for fluorescence observation (see the “Fluorescence observation” section below).

### Cell damage experiments using hydrogen peroxide (H_2_O_2_)

Hydrogen peroxide (FUJIFILM Wako Pure Chemical Corp., 084-07441) was added to NPCs at a final concentration of 250 mM, 500 mM, 750 mM and 1000 mM for 5 min. Then the wells of a 6-well plate were washed three times with the NPC medium, followed by the live and dead cell viability assays and Hoechst imaging (Fig. S2 of the Supplementary Information).

### Phase-contrast microscopy

The human iPS-derived NPCs and neurons were observed by an inverted microscope (IX83, Evident Corporation, Tokyo, Japan), with a 10× phase objective lens (UPlanFLN 10X, Olympus). Images were recorded at 33 frames per second with pixel size of 1.3 µm by a CMOS camera (ORCA-Flash 4.0 V2, Hamamatsu Photonics).

### Fluorescence observation

Fluorescence imaging was carried out on the same microscope that was used for phase-contrast microscopy, equipped with fluorescence filter cubes. A 40× objective lens (UPlanFLN 40×, Olympus) was used to acquire ICC images, and a 10× objective lens (UPlanFLN 10×, Olympus) was used for cell counting. Alexa Fluor 488, Cy3, and Hoechst 34580 signals were visualized using the GFP-3035D (Semrock, Cat# GFP-3035D-000), TRITC-B (Semrock, Cat# TRITC-B-000), and U-FUNA (Olympus, Cat# U-M723) filter sets, respectively. Cells stained with appropriate fluorescent markers were imaged using the corresponding excitation/emission wavelengths.

### Cell counting software

Hoechst-stained fluorescence images were processed and analyzed by using Python (v3.x) with NumPy, SciPy, and OpenCV for automated cell counting based on the multiple ROI scanning method (Fig. 4b); details are provided in Figs. S3-S5 of the Supplementary Information.

### Calculation of errors

For each NB-containing medium (Table1) for Day *N*, each microscopic image covered an area of 1.77 mm^2^, and 10 images were randomly taken to observe the cells plated on the dish with an area of approximately 10 cm^2^. Note that detailed protocols for the cell experiments are described in Fig. S6 of the Supplementary Information), which also indicates that a total of seven independent experiments were performed in this study. *x_i_*(*N*) represents the number of live cells in the *i*-th image (*i*=1,2,3…10). Here, the number of live cells was counted using the self-made Python program (Figs. S3-S5 of the Supplementary Information). The average number *x̅*(*N*) was calculated as:

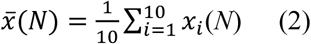

The standard deviation (SD) of *x_i_*(*N*) was computed as:

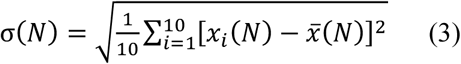

The corresponding average live cell concentration of Day *N* was calculated as:

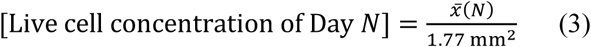

The error of LCR(*N*) in Eq. 1 was calculated using the following standard error propagation formula:

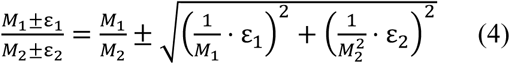

where M_1_= *x̅*(*N*), M_2_= *x̅*(0), *ε*_1_ = *σ*(*N*), and *ε*_2_ = *σ*(0). These analyses were performed using Python (version 3.X) with scipy.stats, numpy, and opencv.

### t-test

Statistical comparisons between groups were performed using two-tailed unpaired Student’s t-tests implemented in scipy.stats (Python). Mean intensity values were significantly different between live and dead cells (Fig. 4c; *t* = –30.613, *p* =7.7 × 10^−68^). Statistical significance was defined as follows: *p* < 0.05, *p* < 0.01, and *p* < 0.001. A *p*-value < 0.001 was considered highly significant.

## Data availability

The data that support the findings of this study are available from the corresponding author upon reasonable request.

## Acknowledgment

We acknowledge Y. Zeng for help in revising Python code for analyzing distributions of zeta potential and size for NB media.

## Funding

This work was supported by JST SPRING, Grant Number JPMJSP2108 to Y.L. M.A.S. was funded by the Natural Sciences and Engineering Council of Canada (NSERC) grants ALLRP 581004-2023 and RGPIN-2024-06341.

## Author contributions

Y.L. performed the experiments and data analysis. T.O. provided NB supplemented culture media with the help of E.K., and associated protocols. M.A.S. and J.S. taught Y.L. and K.H. the cell culture and ICC methods. J.O. developed the basic part of the particle detection program. K.H. designed the experiments together with T.O., and wrote the manuscript in collaboration with L.Y. and T.O. Y.H. participated in discussions related to the research and provided advice on NB research.

## Competing interests

Co-author Dr. Ohdaira is the founder of Ohdaira Laboratory Co., Ltd. (Saitama, Japan), which developed the nanobubble production method used in this study (Japanese Patent Nos. 7637707 and 7227694; US Patent No. 12329152; US Application No. 16/770528; Chinese Patent No. ZL201880088629.2; Korean Patent Nos. 10-2667835 and 10-2657332; European Patent No. 3721886). This relationship has been disclosed to the publisher and does not affect the objectivity or neutrality of the research. All other authors declare no competing interests.

## Notes

### Summary of Updates

In response to the minor revision, we have revised the title and the competing Interests statement accordingly.

