## Supplementary Figures for "Charged nanobubbles in culture media differentially affect viability of human iPSC-derived neurons"

### Development of charged nanobubble-containing media and their effects on iPSC-derived neuron cultures

**Theoretical validation of diffusion for observed nanobubble trajectories.**

To support the size of the nanobubble belongs to nanoscale shown in Fig. S1, we estimated theoretically the diffusion length L:

$$L = \sqrt{2D\tau} \qquad \text{(S1)}$$

where $D$ and $\tau$ are the diffusion coefficient and diffusion time respectively, by using the following Einstein relation;

$$D = \frac{k_B T}{6\pi\eta r} \qquad \text{(S2)}$$

where $k_B$, $T$, $\eta$, $r$ are the Boltzmann constant, temperature, water viscosity and radius of a nanobubble respectively.

Given $\eta$=0.89×10$^{-9}$ pN·s/nm$^2$, a particle radius $r$=500 nm, $k_B \cdot T$=4 pN·nm ($T$ = 25 °C), and diffusion time $\tau$ =10 s, the diffusion coefficient $D$ and theoretical diffusion length $L$ were calculated as:

$$D = \frac{4}{6\pi \times 0.89 \times 10^{-9} \times 500} \approx 477\ \mathrm{nm^2/s}$$

$$L = \sqrt{2 \times 477 \times 10} \approx 97.6\ \mathrm{nm}$$

This result confirms that a nanobubble with a radius of approximately 500 nm would be confined within a nanoscale region (≤ 100 nm) in 10 s, aligning with the representative trajectory shown in Fig. S1. This estimation supports the interpretation that the particle exhibits constrained nanoscale motion consistent with theoretical expectations and validates that its size is below 1000 nm, thereby supporting its identification as a nanobubble.

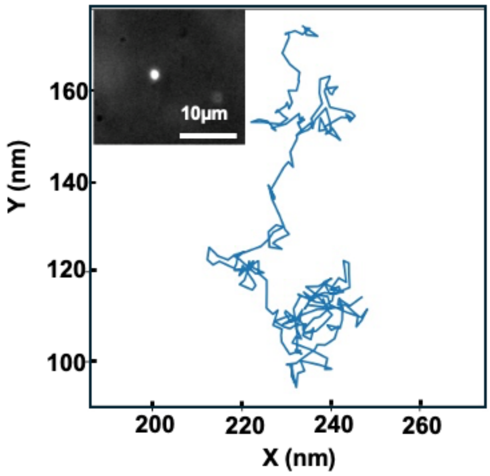


**Fig. S1 The Brownian motion of a single positively charged NB**. The motion of a NB particle, which was generated 43 days prior, was observed by dark-field microscopy at a recording rate of 33 frames per second (Methods). Scale bar: 10 μm. This observation demonstrated that the NBs could remain stable over extended periods. Using our self-developed software (Supplementary Information), its trajectory over 10 seconds was analyzed.

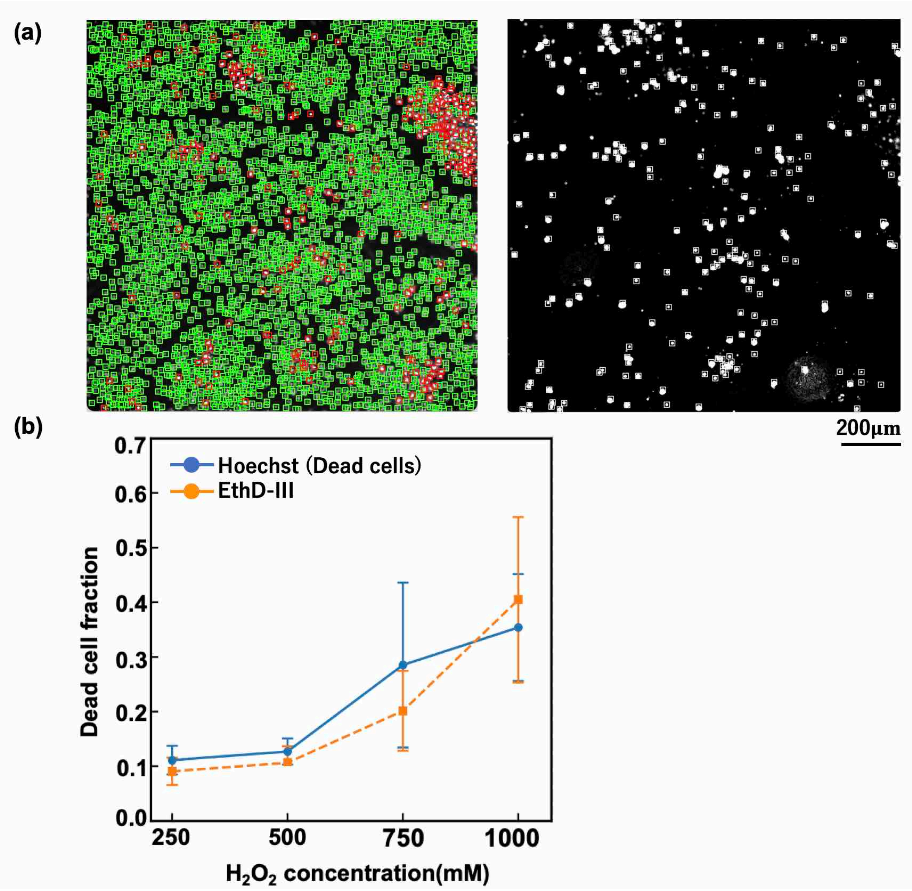


**Fig. S2 Cell damage experiment using hydrogen peroxide. (a)** Example cell-counting results for fluorescence micrographs of the same cells, showing Hoechst-stained nuclei (left) and EthD-III–stained nuclei (right). In the Hoechst imaging micrograph, as shown in Fig. S4, red indicates dead cells, whereas green indicates live cells, as classified according to the criteria shown in Fig. 4(b) of the main text. Because both staining reagents label cell nuclei, cells can be detected using the overlapping ROI scanning method (Figs. S3-S5). Note that the total numbers of the cells were counted from the Hoechst images. **(b)** The number of dead cells was calculated for cells exposed to $H_2O_2$ at concentrations of 250 mM, 500 mM, 750 mM, and 1000 mM for 5 min. For each concentration, measurements were performed in independent wells, and cell counts were calculated from 10 images acquired from a single well. The error-bars represent SD.

**Overlapping ROI scanning method for cell counting**

We developed an overlapping ROI scanning method tailored for a round particle such as a Hoechst-stained nucleus, where its intensity is maximal at the center. Each fluorescence image is scanned with a Region of interest (ROI), whose size is 16 × 16 pixels, corresponding to a nucleus diameter (Fig. 6a of the main text). The ROI is shifted by 1/4 of its size in both x- and y-directions (the black square in Fig. S3a). This overlapping strategy guarantees that every nucleus is captured at least once within the central area of an ROI (Fig. S3c-e). For each ROI, the brightest pixel ($I_{\max}$) and its position ($x_{\max}, y_{\max}$) are obtained, and only ROIs where the position is located within the central half of the ROI are retained as valid detections (Fig. S3b). Duplicate detections of position ($x_{\max}, y_{\max}$) are removed. Based on intensity thresholds, a nucleus is classified into live cells (23 a.u. ≤ $I_{\max}$< 100 a.u.) or a dead cell (100 a.u. ≤ $I_{\max}$ < 230 a.u.). Finally, boxes are drawn on the image to visualize the results (green = live, red = dead) (Fig. S4), and the total counts of live and dead cells are output. This algorithm is summarized in Fig. S5.

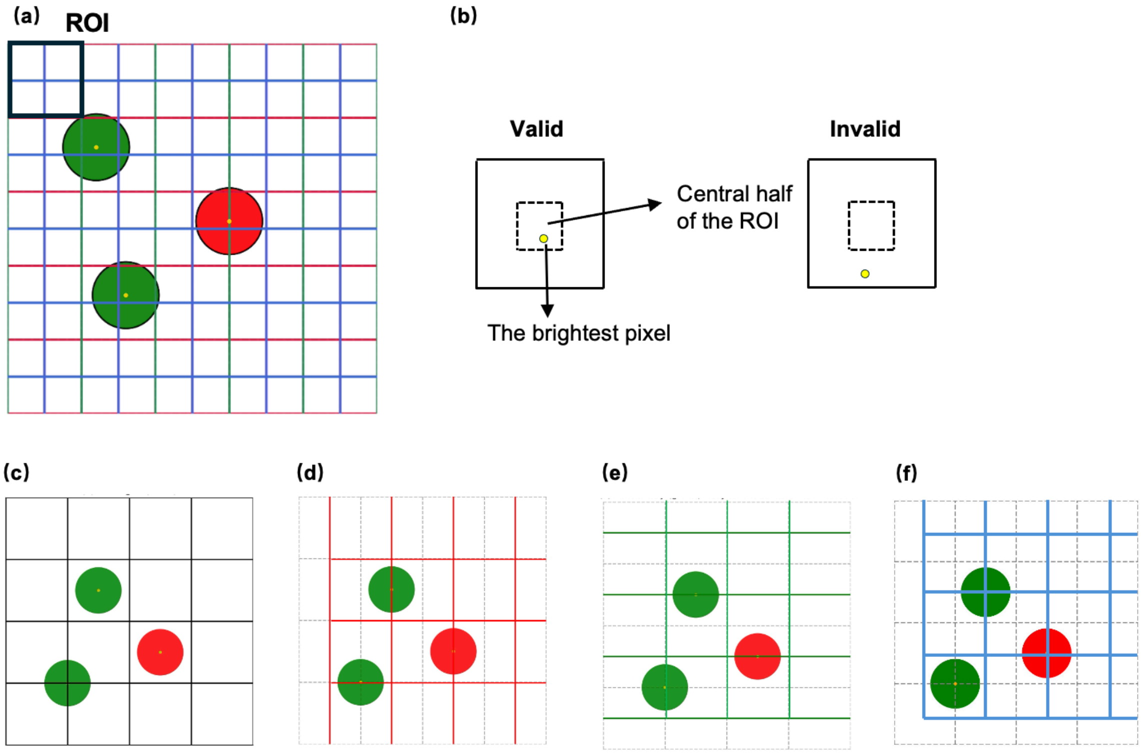


**Fig. S3 Example of the overlapping ROI scanning method in the case that the ROI is shifted by 1/2 ROI size. (a) (b)** Example of valid and invalid detection. **(c–f)** Individual examples of the shifted four grids in Fig. S3a.

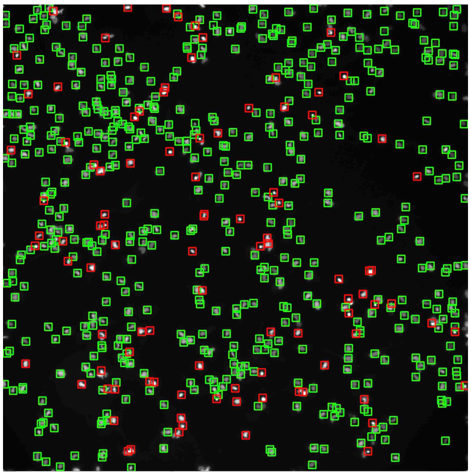

**Fig. S4 Live/Dead cell counting.** Using the FI threshold (Fig. 4b of the main text), both live cells (green) and dead cells (red) can be automatically detected.

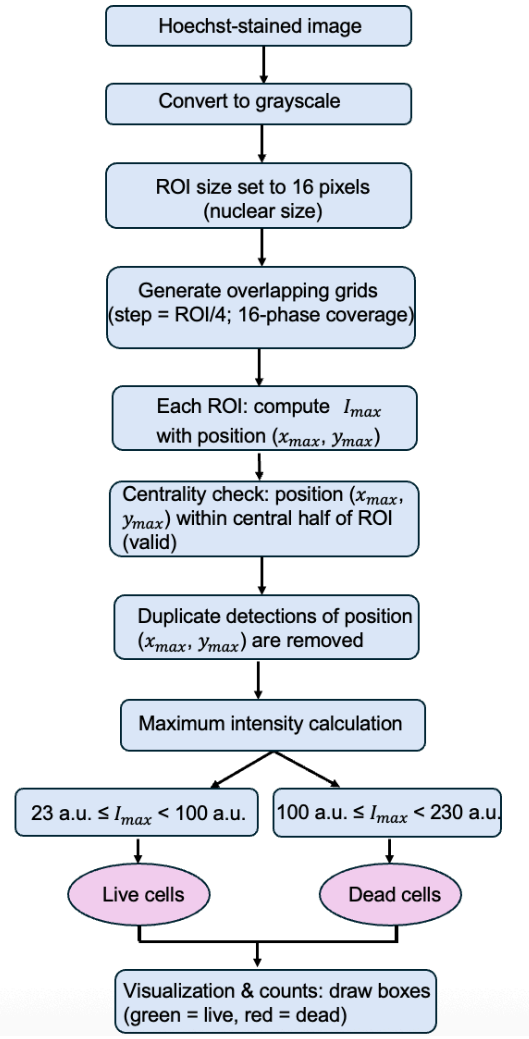


**Fig. S5 Algorithm of custom-built software (the overlapping ROI scanning method for cell counting).**

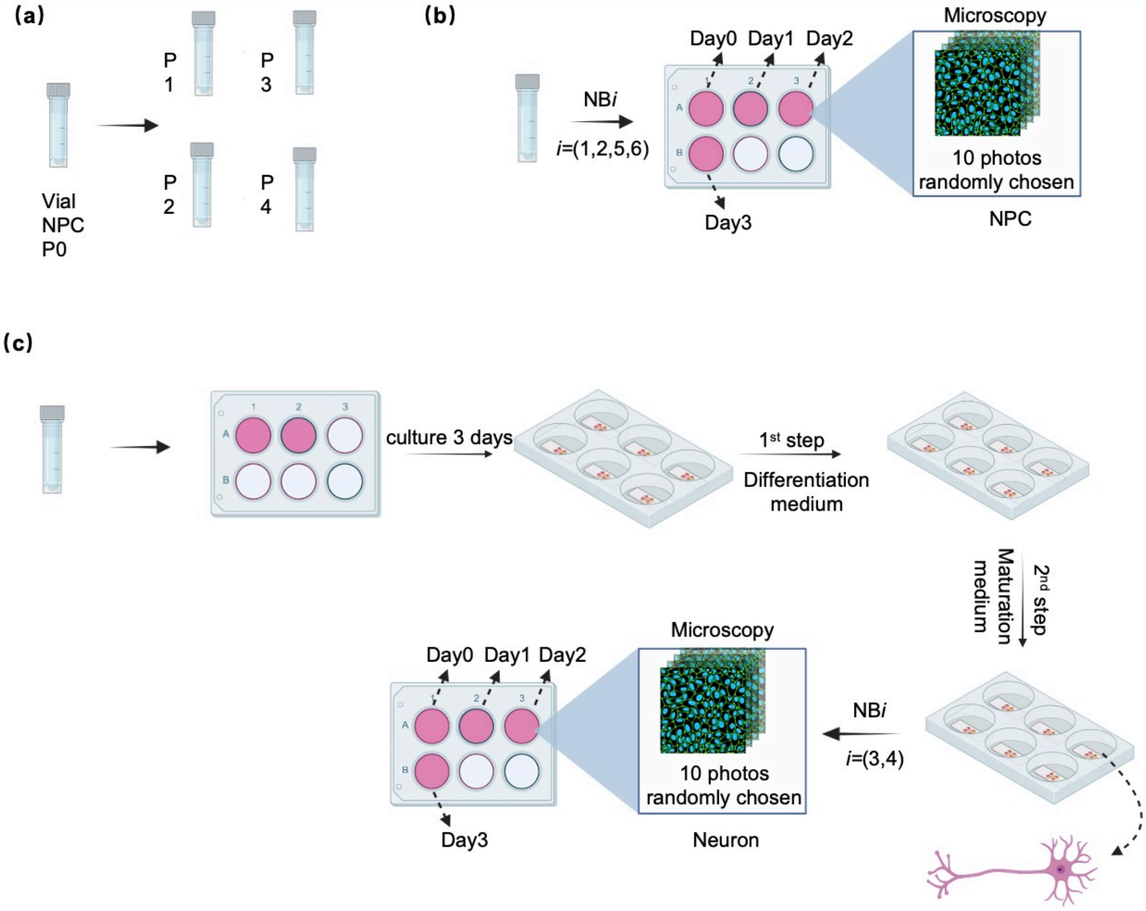


**FIG. S6 Experimental protocols for the cell experiments. (a)** NPCs were expanded from a vial purchased from STEMCELL Technologies, and cells with passage numbers ≤ P4 were used in the experiments. **(b)** In the case of NPCs, for each NB medium listed in Table 1 of the main text, cells expanded from one independent vial were expanded into four wells, which were used for experiments on Day 0, Day 1, Day 2, and Day 3, respectively. Note that after the fluorescence observation, the cells were discarded. **(c)** The protocol for the neurons corresponding to Fig. S6b.

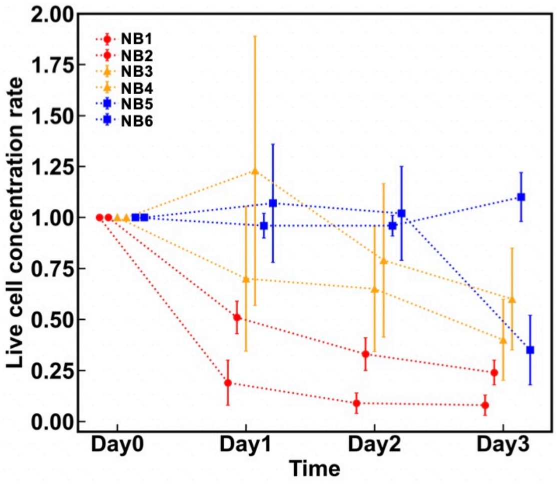


**Fig. S7 Summary of results.** The LCR results (Eq. 1 of the main text) were superposed for all media (NB1–NB6) in Table 1 of the main text. The results for NB4, not shown in the main text, are also presented.
